# Assessing short- and long-term variations in diversity, timing, and body condition of migratory frugivorous birds

**DOI:** 10.1101/2020.10.27.356709

**Authors:** María Campo-Celada, Irene Mendoza, Ana Benítez-López, Carlos Gutiérrez-Expósito, Julio Rabadán-González, Pedro Jordano

## Abstract

Under current global change context, climate change is driving substantial phenological mismatches between plant species and the organisms that rely on them. Given that frugivorous birds are fundamental for forest regeneration, and most of them are migrant species, identifying the effect of global change over them must be a priority. In this study we have analysed changes in the composition, morphometry, and physical condition in an avian community at long- (40 years) and short time (seasonal) spans. Our findings indicate a profound transformation at practically every level of analysis. In 40 years, the avian community shows a 66% and 13% decrease of the wintering and seed-disperser species, respectively. Seasonal abundance peaks were advanced in 13 out of 15 species. In addition, we have found a significant 1.5% increase in the morphometric measurements of certain species, supporting findings in previous studies, and also a remarkable general decrease of body condition. Our results point towards land use changes and climate change as the main causes. If this influence continues to rise, biodiversity will likely be irreversibly altered, damaging crucial ecosystem functions such as animal-mediated seed dispersal and forest regeneration in particular.

## Introduction

Global change drivers are profoundly altering Earth’s biodiversity, ecological systems, and the timing of organisms’ life cycles, i.e., their phenology (Bellard et al., 2014; Hansen et al., 2001; Pachauri et al., 2014). In particular, avian communities are undergoing major changes in their abundance (Inger et al., 2014; Studds et al., 2017), composition, and migratory timing due to land use modifications and alterations of climatic conditions (Fischer et al., 2018; Socolar et al., 2016). These changes result in a widespread decline of several bird species around the globe that, although apparent since the 1980s, has experienced its greatest fall in the last pair of decades (Howard et al., 2020; Wilcove & Terborgh, 1984).

Among bird species, those that undertake migratory movements are under a strong selection because of their site fidelity and fine tuning with available resources. As a result, they are highly susceptible to ecosystem perturbations, compromising returning flights to breeding grounds or resource tracking, for instance (Cuadrado, 1992; Howard et al., 2020; Winger et al., 2019). Thereby, current global change has increased already high migratory pressures over those species, which translates into changes in community composition both in abundance and species turnover over both short- and long-term scales (Howard et al., 2020; Socolar et al., 2016; Wilcove & Terborgh, 1984). Its impact has also proven noticeable in bird species’ morphological trends (e.g., Weeks et al., 2019).

Probably the most overlooked and yet undefined global change-driven effect is related to species phenological mismatches, which occur when there is a temporal uncoupling of interacting organisms. An interesting example is migratory fruit-dependent birds and their food plants (Snow & Snow, 1988). Frugivorous birds rely on fleshy fruits for acquiring the nutritional and energetic resources needed for breeding and migrating (Jordano, 2014). This means the timing of bird arrival or fattening needs to match the timing of fruit crops ripening so that birds can acquire the necessary nutrients for breeding or building up the fat accumulation for migration; meanwhile the fruits are consumed and their seeds effectively dispersed. Yet, climate change is profoundly altering species phenology, resulting in temporal uncouplings. Birds had advanced, for instance, their reproductive cycles and arrival dates when food resources are not at their maximum peak availability, with negative consequences for offspring survival, bird condition and the potential recruitment of biotically-dispersed plant species (see e.g., Nowak et al., 2019; Saino et al., 2011; Visser et al., 1998).

Such temporal uncouplings generate a substantial decrease in the probability of interspecific encounters between interacting partners and may eventually result in the loss of the ecological pairwise interaction. When interactions disappear, the associated ecological functions and services are lost (Valiente-Banuet & Verdú, 2013), with pervasive consequences for ecosystem dynamics (Aamidor et al., 2011; Blendinger et al., 2015; Carnicer et al., 2009; Cooper et al., 2015; Jordano, 1988). For instance, increased warming has substantially advanced the first flowering date of many plant species from the Northern Hemisphere (Menzel et al., 2006), triggering phenological uncouplings with the emergence of pollinator species. Moreover, global changes have impacted on birds’ body condition and morphological traits, as well as phenology and reproductive success (Bailey et al., 2020; Cooper et al., 2015). However, the empirical data on these temporal mismatches and timing shifts in natural processes are very limited.

Migratory birds have been found to show reductions in size over the last decades. For instance, using a four-decade series of specimens from North America, Weeks et al. (2019) showed a consistent decrease in body size across different taxa, coupled with a general increase in birds’ wing length. Body size reductions are likely to lead to a change in shape that compensates for increasing selective pressures in migratory bird species (Weeks et al., 2019; Winkler & Leisler, 1992). The most common shape transformations are elongation of upper limbs, but wing chord and tarsus length decreases have also been reported in other studies, with meaningful differences found between short- and long-distance migratory species (Van Buskirk et al., 2010; Weeks et al., 2019; Winkler & Leisler, 1992). General trends of body change in bird species are difficult to discern, being ultimately related to not only biological factors but also measuring protocols (Labocha & Hayes, 2012). In the short-term, body and fat mass fluctuations that are consistent with long-term reductions have been reported, but they seem to be mainly related to food availability and seasonal weather (Hampe, 2008; Smith, 2016; Tellería et al., 2013; Weeks et al., 2019), which also has a remarkable impact on migration success (Rojas et al., 2019). Both long- and short-term trends in body and fat mass were found consistent with a response to a warmer climate (Gardner et al., 2014; Van Buskirk et al., 2010; Weeks et al., 2019). By finding overall trends across species types, locations, and environmental heterogeneity we could identify generalities that will help forecasting future impacts of global change. We aim to provide empirical evidence to test these predictions by analysing changes (long-term) in wing, tail, tarsus and bill lengths, and changes in fat accumulation (short-term) between decades.

Although fat accumulation is a frequently used proxy for physical condition, morphometric indices of body condition have been found to be more reliable, yet usually well-correlated with fat content (Labocha & Hayes, 2012), or lean dry mass in birds (Schulte-Hostedde et al., 2005). There is no universal-best body condition index, but weight/tarsus residuals based on ordinary least squares regression are independent of the size measurements (predictors; Labocha & Hayes, 2012), and seem to be the best body condition index in our case. Migratory birds’ body condition is determinant on timing, route choice and migration fitness (Bairlein & Gwinner, 1994; Duijns et al., 2017); and body condition relies, for strong frugivorous species, mainly on the availability of appropriate fruit resources (in quantity and quality) to support the energetic demands of migration (Bairlein & Gwinner, 1994; Brown & Sherry, 2006; Parrish, 1997; Stoate & Moreby, 1995). We wanted to check whether body condition-fruit variability matching is different now compared to the 1980s, both at seasonal and interannual scales, given that possible disturbances on fruit production at the stopover area may lead to major ecological consequences.

Although there is ample evidence that birds’ phenology changes as a consequence of direct and indirect effects of climate change, such as food scarcity (especially critical in migratory birds), and mismatches between food peak and breeding periods (Haest et al., 2020; Koleček et al., 2020; van Schaik et al., 1993), few studies have the opportunity to test seasonal variations using a standardized methodology over the long-term. In this study, we focused on frugivorous birds, the long-term changes in community composition and diversity, and their changes over time in morphology and body condition related to fruit availability. We used data on bird abundances and morphometry at two different time scales: short- (with biweekly censuses) and long-term (with historical - 1981-1983 - and current data - 2019-2020; see below). We assessed abundance-based community changes and species turnover in different migratory, trophic, and functional groups. We intended to gain insight in how bird abundances, diversity, phenological timing, body condition and morphology may have been altered in a 38-year period in a Mediterranean sclerophyllous shrubland community. In particular, we aimed to answer the following four specific questions:

1. How has the avian community changed between the 1980s (1980-1983) and 2019-2020 period? What is the extent of turnover for species with different migratory behaviour, trophic level, and ecological function?
2. Is there a seasonal variation in bird abundances?
3. What are the temporal changes in birds’ body size over long- and short-term scales? Are there differences among resident and migrant frugivorous species?
4. How does birds’ physical condition change at seasonal and interannual scales?

## Materials and methods

### Study site

#### Location

The study area was located in Hato Ratón, a locality immersed in Doñana’s Natural Area, close to Villamanrique de la Condesa, Sevilla province, southern Spain (37° 10’ 26.4” N, 6° 20’ 17.4” W, 11 m a.s.l.). Doñana’s Natural Area is an Andalusian protected area of almost 122,500 ha that has been widely studied due to its singular biodiversity. As a natural open space situated at one of Europe’s most southerly points, it is a common breeding and nesting spot for bird species that migrate between Europe and North Africa. Its large abundance of fleshy-fruited shrubs attracts a particularly high number of frugivorous bird species of different migratory behaviours, which makes it a suitable location for our project. The vegetation of the site is characterized by tall, diverse, sclerophyllous shrubland on sandy soils dominated by *Pistacia lentiscus* (Anacardiaceae) and *Olea europaea* var. *sylvestris* (Oleaceae) intermingled with *Pinus pinea* trees (Table S1).

#### Climate

The climate of the study area is typically Mediterranean, with hot and drought periods in the summer that contrast with concentrated rainy months in spring and early autumn. Extreme events are also common in terms of drought or heavy rainfall. Average annual mean temperature and accumulated rainfall in the 1980s were 16.7 °C and 547 mm, respectively, while in the 2010-2019 decade these values were 17.6 °C and 497 mm (Table S2; 1980-2020 series, data from a weather station placed at Doñana Biological Reserve, 20 km from the study area). While mean and minimum temperatures show a slow increasing trend, accumulated rainfall is considerably variable among years and decades. It is remarkable that the pair of years of the first study period (1981-1983) were exceptionally dry within the decade; accumulated rainfall per month barely surpassed the 300 mm. Also, monthly minimum temperatures were colder in 1981-1983 than the whole 80s-90s period.

### Data collection

Data were collected periodically between January 1981 to October 1983, and again between July 2019 to March 2020, which we will refer to these two time periods as the 1980s and 2019-2020 respectively. Data for the 1980s period were previously available, concerning fruiting phenology and fleshy-fruit production, bird community composition, bird morphometry and body condition, diet composition, etc. (Jordano, 1984, 1985, 1987, 1988). Sampling frequency differed between decades, being weekly during the 1980s and biweekly in 2019-2020 (Figure S1).

#### Bird sampling

Between 6 and 10 nets were operated weekly during 1981-1983 and 12-20 biweekly in 2019-2020. Mist nets were opened from dawn to dusk (1981-1983) or midday (2019-2020) and checked at hourly intervals. Trapped birds were collected individually in fabric bags and then identified and measured by expert ringers following the standard procedures of the EURING criteria. Individual birds were ringed, and the following variables were recorded: species, age, sex, type of moult, existence of brood patch, fat and muscle and also measured body mass (±0. 5 g) and wing, F8 feather, tail, tarsus, and bill (culmen) lengths (± 0.1 mm; ± 0.5 mm for wing, F8 feather, and tail). We checked for data repeatability to assure that measurements between different ringers were significantly consistent. We obtained a significantly high correlation between measures performed by different ringers (r = 0.92-0.99 for all measurement variables, Figure S2), and a significantly high measurement repeatability (i.e., intraclass correlation coefficient, R package “rptR”, Stoffel et al. 2017; N= 5000 resamplings) for most linear measurements of species with sufficient sample size (e.g., *S. atricapilla*, *E. rubecula*, *S. melanocephala*) (R > 0.793, p << 0.01 for all variables except tail length in *E. rubecula* with p < 0.08).

We captured 2,257 birds in 1981-1983 and 780 in 2019-2020 (more than 3,000 individuals in total), corresponding to 60 species with different migratory strategies, diet, and functional role. To have sufficient sample sizes for the morphological and body condition analysis, we selected only the 6 species for which we had more than 20 complete captures (including collected data for all study variables) for each period of time: 1981-1983 and 2019-2020. Those species are Eurasian blackcap (*Sylvia atricapilla*), Sardinian warbler (*Curruca melanocephala*), common blackbird (*Turdus merula*), European robin (*Erithacus rubecula*), European greenfinch (*Chloris chloris*) and common chiffchaff (*Phylloscopus collybita*). All recorded species were named following The IOC World Bird List taxonomy updates (Gill, Donsker & Rasmussen, 2020), completed with Spanish common names included in The List of Birds of Spain, 2019 (Rouco et al., 2019).

Mist-netting was complemented with bird censuses using linear transects, recording each individual that was seen or heard, together with information on the species and hour, following the methodology used in 1981-1983 (Jordano, 1984a, 1985). Censuses were completed at least twice a month along a 2 km-line transect located across the study area. All censuses started within one hour after sunrise and were performed avoiding extreme weather conditions.

#### Fruiting phenology

Seasonal and annual fruit abundance was quantified by counting all fruits (ripe and unripe) present in linear transects. In 1981-1983 a total area of 187.5 m^2^ was sampled, whilst in 2019-2020, the total area was 1,350 m^2^. Main species included *Pistacia lentiscus*, *Rhamnus lycioides, Phillyrea angustifolia*, *Olea europaea* var. *sylvestris*, *Ruscus aculeatus*, *Pyrus bourgaeana* and *Myrtus communis,* among others. Fruit production was determined when fruits were unripe and had reached their final size, before the ripening and consumption time. Phenological variations were also monitored weekly in 1981-1983 and biweekly in 2019-2020, up to 15 and 26 sampled days, respectively. Total fruit production was 7,596,942 fruits/ha in 1981-1983, and 49,708.29 fruits/ha in 2019-2020 (152 times higher in 1981-1983) in our study area.

In 1981-1983, we recorded 92.1 % of area covered by woody vegetation, and 7.9 % of bare ground (Table S2). Up to 16 species of fleshy-fruited shrubs and treelets were recorded and monitored in the area, representing 72.1 % of the vegetation cover, which was dominated by the fleshy fruited species *Pistacia lentiscus* and *Smilax aspera*, followed by leguminous *Ulex parviflorus*. On the other hand, we detected a marked vegetation transformation in the 2019-2020 period. Although the percentage of bare ground had increased up to 27.5 % (against 72.5 % of area covered by vegetation), we observed a considerable vegetation shift in our study plots, which became encroached by arboreal species such as *Pinus pinea,* showing a more complex vegetation structure with higher canopy. Fifteen fleshy fruit species were monitored in 2019-2020, accounting for 50.5 % of the vegetation cover, which was dominated by *P. lentiscus* (26.7 %) and the coniferous (non-fleshy fruited) species *Pinus pinea* which increased in our study area from less than 1 % in the 1980s to 21.4 % currently.

### Data analyses

#### Community composition

We classified species according to three criteria: 1) migration type, 2) diet, and 3) ecological function. Regarding migration types, species were distinguished depending on their timing of arrival and departure into four categories: **“Wintering”** (species that spend October-March in the study area), **“Summering”** (species that spend May-September in the study area, where they breed), **“Transient”** (species that spend only a brief period of time, e.g., trans-Saharan migrants using the area as a stopover site) or **“Resident”** (present in the area the whole year); the first three categories belong to migratory species. We also classified species according to their trophic level in two broad groups: **“Frugivores”** sensu lato (including all species which, at least occasionally, feed on fleshy-fruits, e.g. frugivore-insectivore or granivore-frugivore, carnivore-frugivore and omnivore species) and **“Non frugivores”** sensu stricto (the rest of species, with no recorded fruit feeding behaviour) (Jordano, 1987; Snow & Snow, 1988). We further classified species based on their functional role as **“Seed disperser”** (species that contribute seed dispersal events, frequently by regurgitation or defecation of ingested fruits, Snow & Snow, 1988), **“Seed predator”** (species that feed on or damage the fruit seeds), **“Pulp consumer”** (species that feed on fruit pulp and do not contribute to seed dispersal) and **“Non frugivore”** for the rest of species that do not feed on plant fleshy-fruits. Some species switched their behaviour depending on the plant species or the season (“Seed disperser/Pulp consumer” and “Seed disperser/Seed predator/Pulp consumer”, classified as “Seed disperser” for analyses; and “Seed predator/Pulp consumer”, classified as “Non seed disperser” (see Snow & Snow, 1988 for a thorough description). We used the three different classifications (migration type, trophic type, and functional type) to assess changes in the avian community between the two study periods (Table S3).

We calculated species relative abundance (ind/km) per month and study period. As the sample size for some species was limited, we selected the 20 most abundant species per time period to characterize community composition change between decades and seasonal variation in species’ abundances within study periods (Table S3). We compared bird species’ abundance and calculated species turnover between study periods for all species and for species with different migratory behaviour, trophic level, or functional role. We also calculated the proportion of gained and lost species between study periods (1981-1983 & 2019-2020). Species turnover was calculated for each migratory, trophic, and functional category that included at least 5 species in both time points.

#### Migration phenology

Species’ seasonal abundance (monthly mean individuals per km^2^) distribution was used to detect peaks (maximum density) and the date of the first and last recordings per species and year. We used the most abundant species per study period to explore its seasonal variation, and assessed differences between resident and wintering bird species, and between frugivorous and non-frugivorous birds (23 spp. finally). We corrected bird abundances by the number of sampled days per period (50 days in 1981-1983 and 18 days in 2019-2020). We did not sample in April to June in 2020, so these months were removed from the comparison. We assessed the consistency in temporal shifts in abundance peaks between periods by comparing the proportion of species showing advanced shifts in their date of maximum abundance using a binomial test, determining whether temporal shifts to earlier dates occurred more often than randomly expected. To determine total monthly differences per migratory (resident and wintering) and trophic types (frugivores and non-frugivores), we performed four Mann-Whitney tests, each with monthly abundances of resident, wintering, frugivore and non-frugivore species, using both study periods as pairing variables.

#### Morphometry and fat change

For each of the six species with more than 20 captures per decade, we analysed changes in four morphometric variables: wing, tail, tarsus, and bill lengths. ANOVAs were built using morphometric measures as dependent variables, and study period (1981-1983 or 2019-2020), migration type (resident or wintering), species, sex, and their respective interactions as explanatory variables. We performed model selection and selected the best model based on the Akaike Information Criterion (AIC). To determine which species and body measurements were truly different among decades, the direction of the changes and its magnitude, we calculated mean body measure length and its standard error per time period in each of the 6 species. For further detail, we calculated each species’ percentage of increase (or decrease when negative values) for wing, tail, tarsus and bill measurements in 2019-2020 relative to 1981-1983, and then performed their respective species individual ANOVA analysis to test for significance of changes in each case.

We also tested short term changes in fat accumulation for all the captured birds (2,805 individuals). We were interested in intra-annual species fat change evolution between three key periods along the migratory cycle (Gordo, 2007): **“Migratory pass”**, from July to September, when most of our migratory species make a stopover in southern Spain during across-continents routes; **“Autumn step”**, October and November, stopover step before species arrival to wintering destinations; and **“Wintering”**, from December to March, when just wintering and resident species remain in the study area. Fat accumulation was noted with a 0-4 scale in 1981-1983 and a 0-8 scale in 2019-2020 (EURING protocol), thus we standardized both to the same scale by establishing qualitative categories instead: Low fat (0-1 and 0-2), Medium fat (2-3 and 3-5) and High fat accumulation (3-4 and 6-8 points). As we only had a few records of high fat individuals in 1981-1983, we finally collapsed categories into **Low fat** and **Medium-High fat** accumulation, encompassing the EURING categories of 0-3 and > 3, respectively. We performed a Generalized Linear Model (GLM) using fat accumulation categories as response variable, year group and migratory seasons as explanatory variables (also including its interaction), and binomial error as link function. Finally, to visualize fat accumulation levels’ distribution along seasons, we corrected its frequencies per total monthly recordings to equate time series data according to different sample sizes.

#### Body condition

We calculated body condition using residuals of body weight regressed on tarsus length (Labocha & Hayes, 2012). Positive residuals indicate that the birds’ weight is higher than expected according to its size (“good” body condition), whereas negative residuals indicate the opposite (“bad” body condition). We tested whether body condition changed between study periods and for migratory vs resident species and their interactions with a GLM fitted with a Gaussian error. Also, we assessed monthly variations in bird condition in relation with fruit availability. Finally, we separately tested the increase/decrease in body condition in bird species for which we had at least 10 residuals in 1981-1983 and 2019-2020 respectively (Figure 5.C).

All analyses were performed in R Statistical Software v. 3.6.2 (R Core Team, 2019). Repeatability coefficients were calculated using the “rptR” package (Stoffel et al., 2017). Species turnover was calculated using the “turnover” function included in the “codyn” package (Hallet et al., 2020). The selection of the ANOVAs models was based on the “aictab” function included in “AICcmodavg” package (Mazerolle, 2020).

## Results

### Community composition, bird abundance and species turnover across periods

We found a species beta-turnover of the 20 most abundant species of 0.52, with the same proportion (0.26) of species losses and gains between 1981-1983 and 2019-2020 (Table 1). This means that half of species per decade had reduced their relative abundance and disappeared from the 1981-1983 “top 20” list, being replaced in 2019-2020 by new birds that were not present or were scarce in 1980s records (Figure S4). We found that the number and abundance of resident bird species have increased, with a 0.55 beta-turnover between study periods. The proportion of gained and lost species was 0.35 and 0.20, respectively, and the relative abundance of resident species increased from 60% to 80%, approximately (Figure 1.A). In turn, we found that the relative abundance of the four main wintering bird species has fallen almost two thirds (66%) between 1981-1983 and the current period (Table 1 and Figure 1.A). The abundance proportions of frugivores and non-frugivores showed practically no change between study periods (Figure 1.B), but both groups had gone through a considerable species turnover, especially among the non-frugivores (with a turnover value of 0.60, versus 0.47 in frugivores). On one hand, the fruit-eating species as a group maintained its relative abundance, whilst the proportion of new species was lower than the proportion of lost species (0.18 against 0.29). On the other hand, non-frugivore species (granivore, insectivore, granivore-insectivore, and herbivore species) also maintained their abundance but gained the double of species they lost from 1981-1983 to present (0.40 against 0.20).

**Table 1:** Species turnover values of different categories (“Groups”) classified according three criteria (“types”). The number of species included in each group is indicated in column “n”. Species’ groups turnover values (“Turnover”) were also split into proportions of species that appeared (“Gained spp.”) and disappeared (“Lost spp.”) in the 2019-2020 period compared to 1981-1983 study years. Observations were also included in the last column. *: Transient and summering species were excluded from migratory category because they were not sampled in 2019-2020. Wintering species were abundant but did not reach 5 spp. per study period needed to calculate turnover.

| Species turnover |  |  |  |  |  |  |
| --- | --- | --- | --- | --- | --- | --- |
| Type | Group | n | Turnover | Gained spp. | Lost spp. | Observations |
| - | All | 27 | 0.52 | 0.26 | 0.26 | - |
| Migratory behaviour * | Resident | 20 | 0.55 | 0.35 | 0.20 | - |
|  | Wintering | 4 | 0.00 | 0.00 | 0.00 | Density fell 66% |
| Functional | Seed dispersers | 17 | 0.53 | 0.20 | 0.33 | - |
|  | Non-dispersers | 10 | 0.60 | 0.40 | 0.20 | - |
| Trophic | Frugivores | 17 | 0.47 | 0.18 | 0.29 | - |
|  | Non-frugivores | 10 | 0.60 | 0.40 | 0.20 | - |

**Figure 1:**
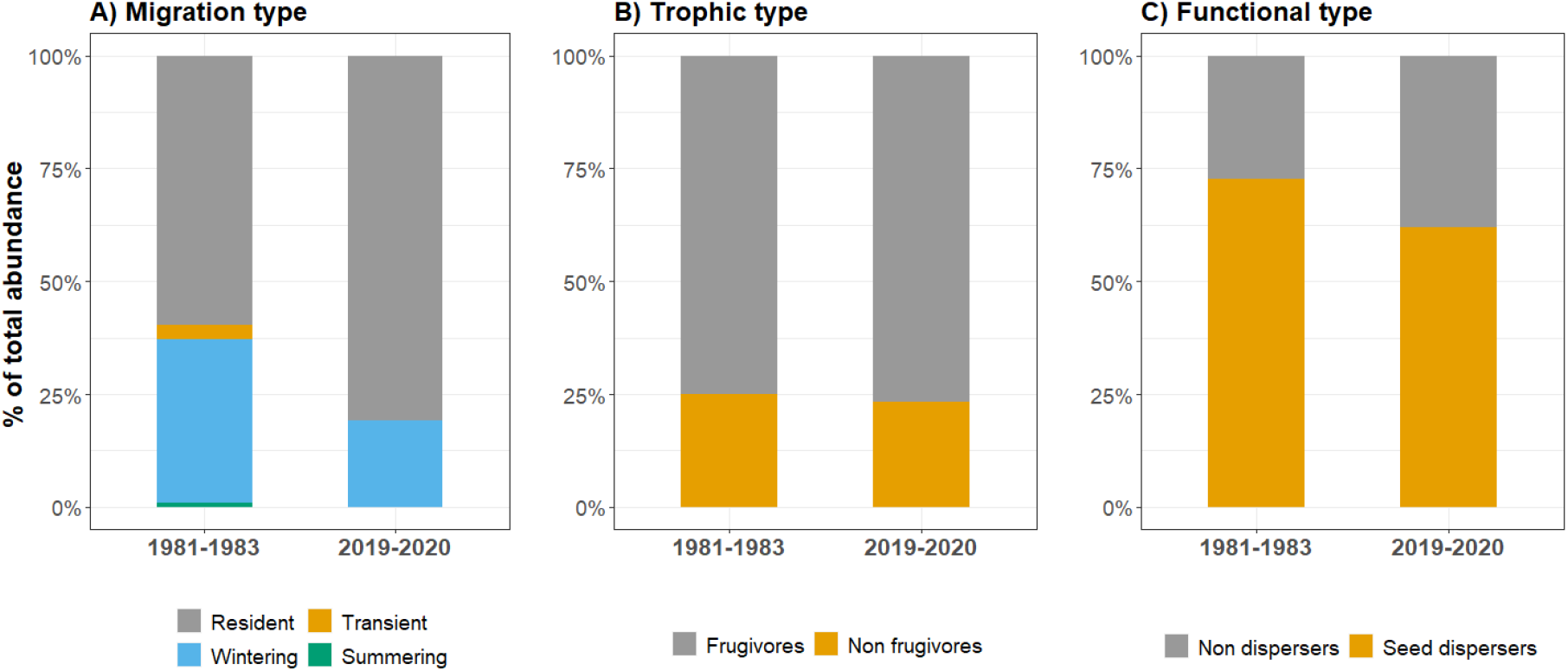
Community composition bar plots showing the percentage of total abundance of the 20 most abundant species during each study period (1981-1983 and 2019-2020), classified according to three different categories: migration (A), trophic (B) and functional types. In total, 27 different bird species were included.

Regarding species’ functional groups, birds acting as legitimate seed dispersers have also experienced a decrease in their relative abundance from almost 75% to 62% (see Figure 1.C), with a beta-turnover value of 0.53, and a negative balance of gained versus lost species (0.22 and 0.33). In contrast, non-seed dispersers have become more abundant, with a high turnover value of 0.60 that reflects twice as much gained as lost species (0.40 and 0.20).

### Seasonal variation in bird abundances

From the 23 species analysed, 8 of them were solely recorded in one study period (7 new species in 2019-2020, and one disappeared; Figure 2). Within the remaining 15 species, we identify peak advancement shifts in 13 of them (10 resident and 3 wintering species), one single delay in wintering *P. collybita*, and no apparent change in *F. coelebs*. A binomial test using the 13 spp. over 15 indicates a consistent, highly significant (p-value: 0.0074), trend across species towards earlier phenologies in the area.

**Figure 2:**
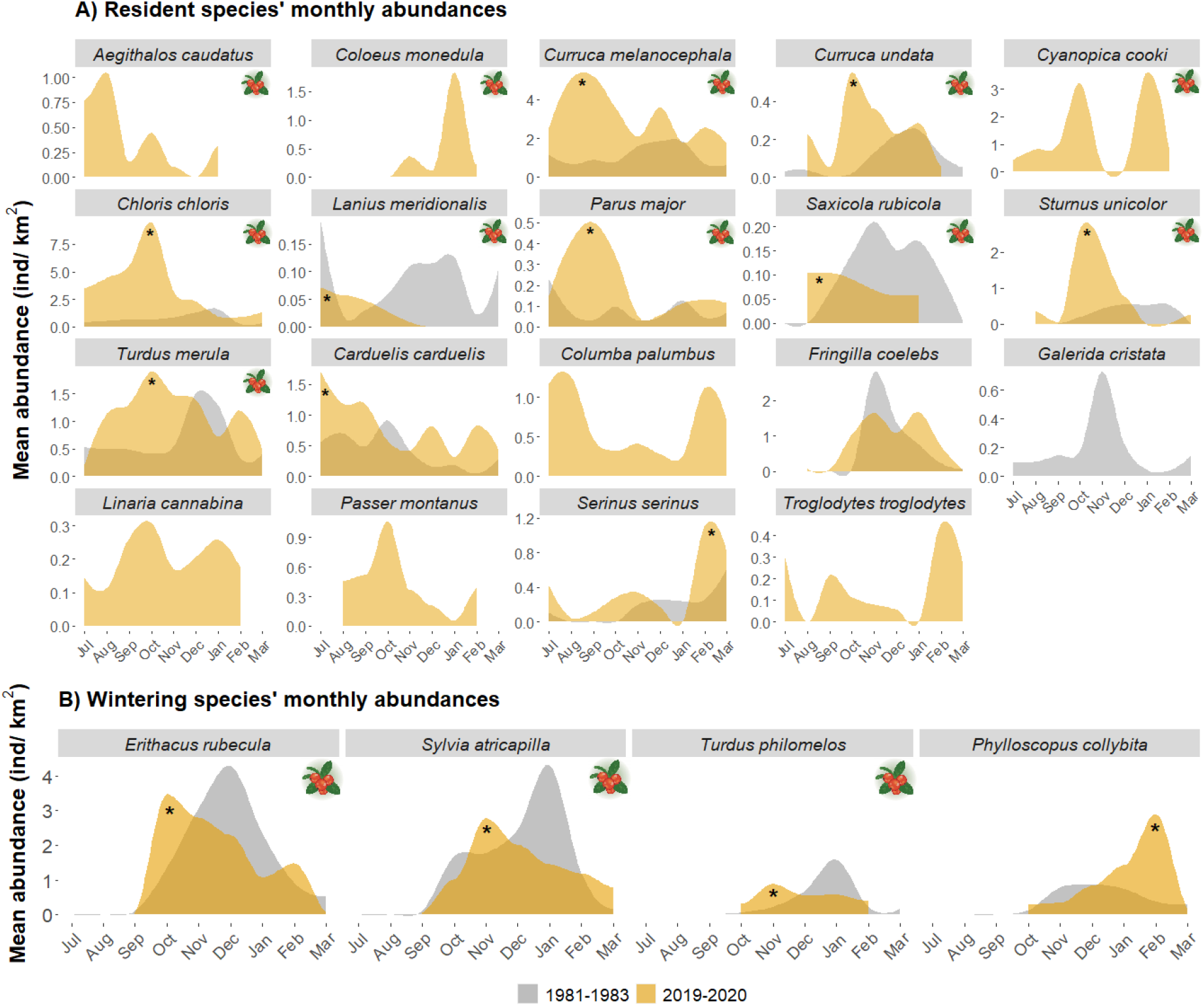
Monthly densities, expressed as individuals per km^2^, of the different resident (A) and wintering (B) species registered in study censuses. Species’ monthly densities of 1981-1983 period are plotted in grey colour, and 2019-2020 recordings appear in orange colour. Symbol with a fleshy fruit means the species is frugivorous. Black asterisks mark the main abundance peak in the 2019-2020 period.

Considering the 23 species trophic types (Table S3), 14 of them have, at least partially, a frugivorous diet and 9 of them never or hardly ever feed on plant fruits. We identified advancements of the maximum abundance date in 11 frugivorous species: *C. melanocephala, C. undata, C. chloris, E. rubecula, L.meridionalis, P. major, S. atricapilla, S. rubicola, S. unicolor*, *T. merula* and *T. philomelos;* and in *C. carduelis* among the non-frugivorous species. On the other hand, the only phase delay belongs to the largely insectivorous *P. collybita* (Common Chiffchaff), which also experienced a considerable increase in abundance in January-February. The other species whose increase deserves to be mentioned are *C. melanocephala*, *C. chloris*, *P. major* and *S. unicolor*, almost doubling their abundances in peak months (Figure 2).

With respect to functional types, 13 out of the 14 mentioned frugivorous species are also legitimate seed dispersers, except *C. chloris*, which is a seed predator (Table S3). Therefore, their phenological patterns are practically identical to those previously mentioned: 13 seed disperser bird species from which 10 show a marked advance in the peak abundance date (Figure 2).

Moreover, when grouping all recorded species’ phenological data into migratory categories (resident, wintering, summering and transient) we observed an overall peak advancement in both resident and wintering species (Figure S3): as we can infer from species’ specific phenological analyses, the general peak advancement in both resident and wintering species is also noticeable in the collapsed resident and wintering groups, respectively. The summering and transient species’ groups were not large enough to elucidate consistent changes. Mann-Whitney tests revealed that monthly abundances significantly differed between study periods in resident (p<< 0.01), wintering (p < 0.05), frugivore (p << 0.01) and non-frugivore (p << 0.01) species groups.

### Temporal changes in birds’ body size at long- and short-term scales

All models including morphometric variables showed lower AIC values when including study period, migrant category, species, and interactions as parameters, although sex was only included for wing and tail length (Table 2). The ANOVA’s tests results showed a significant difference in the four body components that can be explained by study period, migrant categories, species and, except in tarsus length, also by sex variables. We also found significant interactions among study periods and migrant categories in tarsus length, and among study periods and recorded species in bill length.

**Table 2:** Morphometric ANOVAs’ results. Significant values are red coloured. The significance codes are: 0 ‘***’ 0.001 ‘**’ 0.01 ‘*’ 0.05 ‘.’ 0.1 ‘ ’ 1.

|  |  | Df | F value | Pr (>F) |
| --- | --- | --- | --- | --- |
| **Wing** | Year group | 1 | 1447.11 | < 2.2e-16 *** |
|  | Migrant | 1 | 829.03 | < 2.2e-16 *** |
|  | Species | 4 | 12675.40 | < 2.2e-16 *** |
|  | Sex | 2 | 17.07 | 5.28e-08 *** |
|  | Year group:Migrant | 1 | 1.38 | 0.24 |
|  | Year group:Species | 4 | 2.40 | 0.0485 * |
| **Tail** | Year group | 1 | 266.65 | < 2.2e-16 *** |
|  | Migrant | 1 | 1143.33 | < 2.2e-16 *** |
|  | Species | 4 | 3249.21 | < 2.2e-16 *** |
|  | Sex | 2 | 10.15 | 4.38e-05 *** |
|  | Year group:Migrant | 1 | 0.004 | 0.95 |
|  | Year group:Species | 4 | 2.17 | 0.07 . |
| **Tarsus** | Year group | 1 | 398.18 | < 2.2e-16 *** |
|  | Migrant | 1 | 17.81 | 2.69e-05 *** |
|  | Species | 4 | 6934.32 | < 2.2e-16 *** |
|  | Year group:Migrant | 1 | 4.40 | 0.04 * |
| **Bill** | Year group | 1 | 1022.29 | < 2.2e-16 *** |
|  | Migrant | 1 | 1959.49 | < 2.2e-16 *** |
|  | Species | 4 | 4098.88 | < 2.2e-16 *** |
|  | Sex | 2 | 3.41 | 0.03 * |
|  | Year group:Migrant | 1 | 0.13 | 0.72 |
|  | Year group:Species | 4 | 2.10 | 0.08 . |

**Table 3:** Species’ percentage of body component length increase in 2019-2020 compared to the 1981-1983 ones. “Type” column indicates whether species are wintering (“W”) or resident (“R). Note that all species but *Phylloscopus collybita* are frugivores. Negative numbers mean a decrease in body part’s length. Values red coloured were found statistically significant after ANOVA’s. The significance codes are: 0 ‘***’ 0.001 ‘**’ 0.01 ‘*’ 0.05 ‘.’ 0.1 ‘ ’ 1.

| Species | Type | n | Wing | Tail | Tarsus | Bill |
| --- | --- | --- | --- | --- | --- | --- |
| *Sylvia atricapilla* | W | 341 | 1.18** | 1.46** | 0.05 | -0.53 |
| *Curruca melanocephala* | R | 232 | 0.33 | 0.67 | 0.64 | 0.97 |
| *Erithacus rubecula* | W | 118 | -0.27 | 1.9* | -0.64 | -0.34 |
| *Chloris chloris* | R | 117 | 0.71 | 0.84 | 1.4* | -1.27 |
| *Turdus merula* | R | 84 | -1.24 | 3.14 | 1.37 . | -1.73 |
| *Phylloscopus collybita* | W | 37 | -0.04 | 2.42 | 0.55 | -1.7 |

We found a significant difference among resident and wintering species in wing, tail, and bill lengths, but not in tarsus length. Regarding species-specific changes in morphometry, we only found significant differences in three species (Figure 3). Tail length has significantly increased over time for two wintering species: *S. atricapilla* (+1.46 %) and *E. rubecula* (+1.9 %), whereas wing length has increased significantly for *S. atricapilla* only (+1.18 %). Finally, tarsus length has significantly increased in resident *C. chloris* (+1.4 %). Unequal sample size among the two study time periods might be hindering significance for the tail length increase in n *T. merula* or bill decrease found in *P. collybita* case.

**Figure 3:**
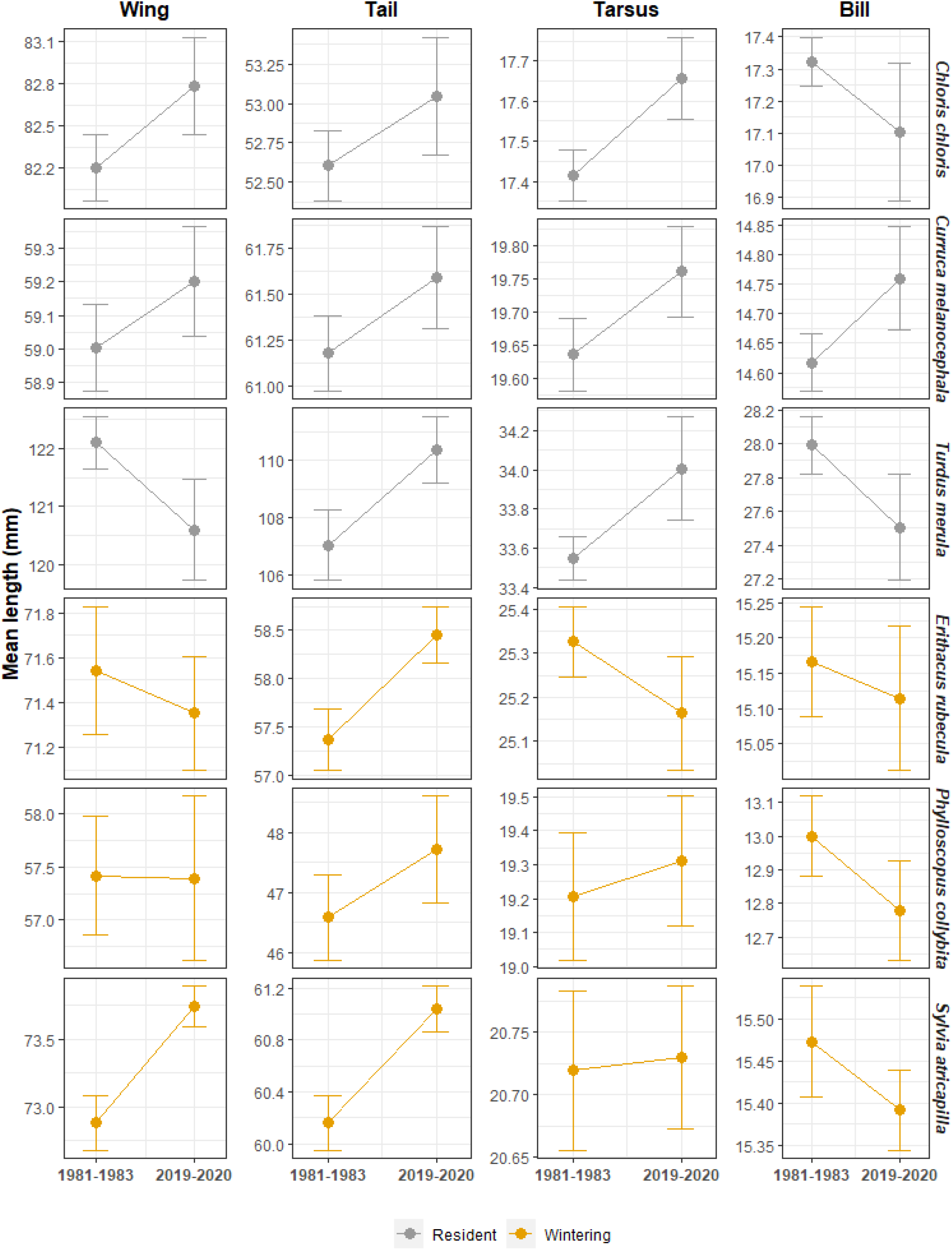
Comparison of wing, tail, tarsus, and bill mean lengths (in millimetres) between study periods (1981-1983 and 2019-2020) for the 6 most abundant species. In grey colour, resident bird species; in orange, the wintering ones.

The GLM’s results show that birds’ body fat accumulation varied significantly between study periods and within migratory seasons (Table 4.A). Birds with a low accumulation of fat in their bodies represented more than 65% of captured individuals in both decades, whilst birds with medium or high fat accumulation accounted for less than one-third of captures in each period respectively (Figure 4). We found two different patterns among year groups: in 1981-1983, medium-high fat accumulation peaked at the migratory pass, while in 2019-2020 it occurred at the autumn step. Low fat accumulation individuals seem to be more, or at least equally, common among migratory seasons, while birds with medium-high fat have been less detected in 2019-2020, especially during the migratory pass season.

**Table 4:** A) Generalized Linear Model (GLM) using binomial error as link function, qualitative fat categories (Low fat and High-Medium fat) as response variable and year group (1981-1983 or 2019-2020) and migratory season (Migratory pass, Autumn step and Wintering) as explanatory variables. B) GLM using Gaussian as link function, weight/tarsus lengths residuals as response variable, and year group and migratory category (resident, wintering, summering and transient) as explanatory variables. Significant p-values are coloured in red. The significance codes are: 0 ‘***’ 0.001 ‘**’ 0.01 ‘*’ 0.05 ‘.’ 0.1 ‘ ’ 1.

| **A) Binomial GLM** |  |  |  |  |
| --- | --- | --- | --- | --- |
| **Response: fat accumulation** | Estimate | Std. Error | z value | Pr (> \|z\|) |
| (Intercept) | -0.43 | 0.09 | -4.90 | 9.84e-07 *** |
| Year group | -1.44 | 0.29 | -4.96 | 7.10e-07 *** |
| Autumn step | -1.20 | 0.13 | -9.26 | < 2e-16 *** |
| Wintering | -1.28 | 0.14 | -9.20 | < 2e-16 *** |
| Year group : Autumn step | 1.94 | 0.33 | 5.88 | 4.05e-09 *** |
| Year group : Wintering | 1.34 | 0.35 | 3.79 | 0.00015 *** |
| **B) Gaussian GLM** |  |  |  |  |
| **Response: weight/tarsus residuals** | Estimate | Std. Error | t value | Pr (> \|t\|) |
| (Intercept) | 8.18 | 0.56 | 14.56 | < 2e-16 *** |
| Year group | -6.19 | 0.88 | -7.07 | 2.44e-12 *** |
| Migratory category | -11.22 | 0.74 | -15.06 | < 2e-16 *** |
| Yeargroup : Migratory category | 4.46 | 1.13 | 3.94 | 8.40e-05 *** |

**Figure 4:**
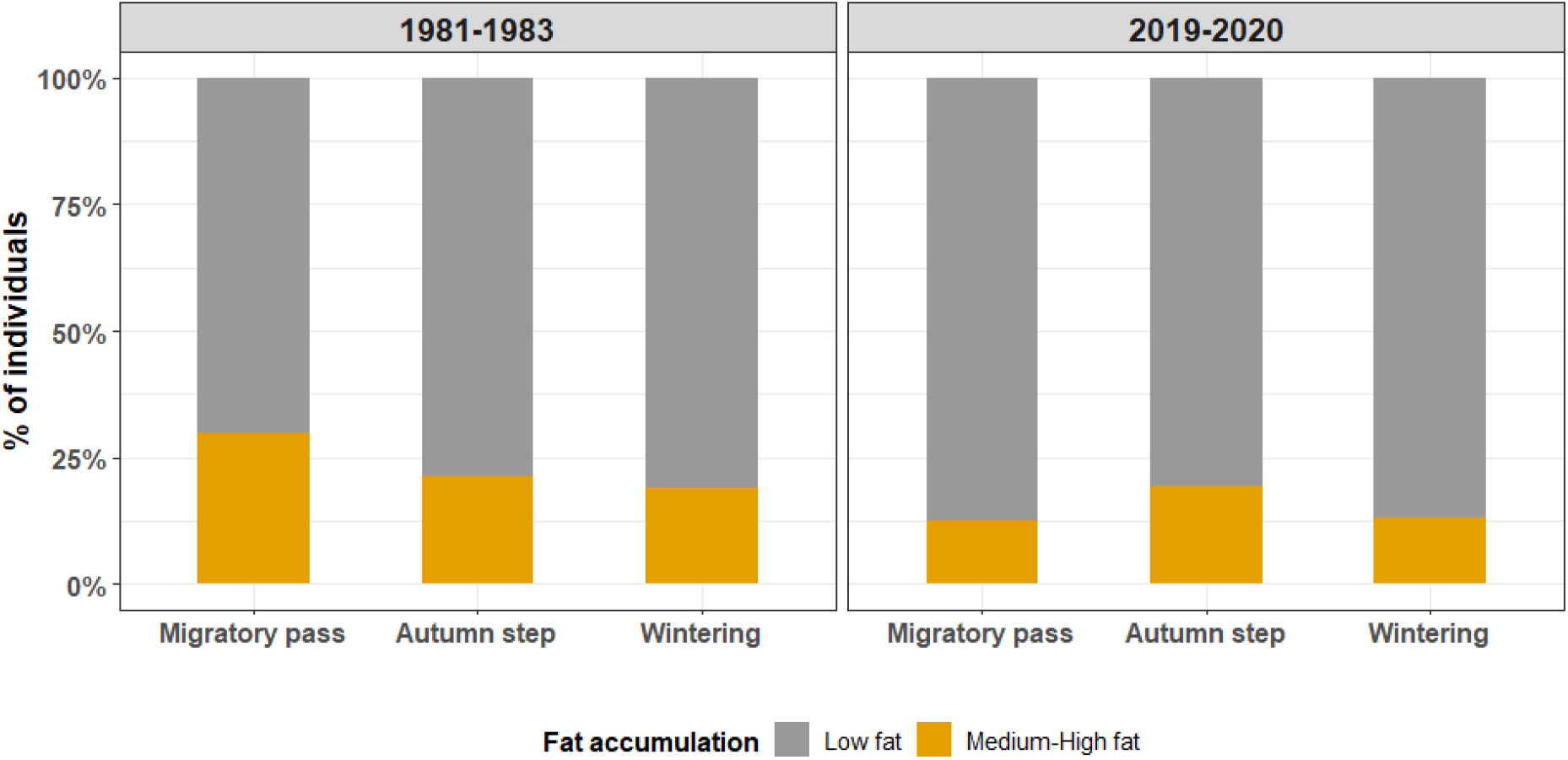
Comparison of seasonal birds’ fat accumulation between study periods (panels “1981-1983” and “2019-2020”), expressed as the percentage of recorded individuals belonging to low fat accumulation (grey colour) or high-medium fat accumulation (orange colour) levels in each migratory season: migratory pass (July-September), autumn step (October-November) and wintering (December-March).

### Change in body condition at seasonal and interannual scale

We found significant differences in body condition among study periods and migratory categories, as well as statistically significant interactions between both factors (Table 4.B). Generally, we recorded more individuals with a good body condition in 1981-1983 than in 2019-2020, with a higher median value in Figure 5.A. Over a seasonal scale (Figure 5.B), we also obtained a trend for higher median body condition monthly value in 8 out of 9 study months when comparing 1981-1983 vs. 2019-2020; pointing to a consistent trend towards reduced general body condition along the year in Hato Raton’s 2019-2020 avian community. At the species level, we identified 7 out of 8 species that presented a relative decrease of mean body condition, although the change was more pronounced for *T. merula* (Figure 5.C). Only *Erithacus rubecula* showed better body condition in 2019-2020 compared with 1981-1983, meaning that a higher proportion of residuals were above 0 in this particular species.

**Figure 5:**
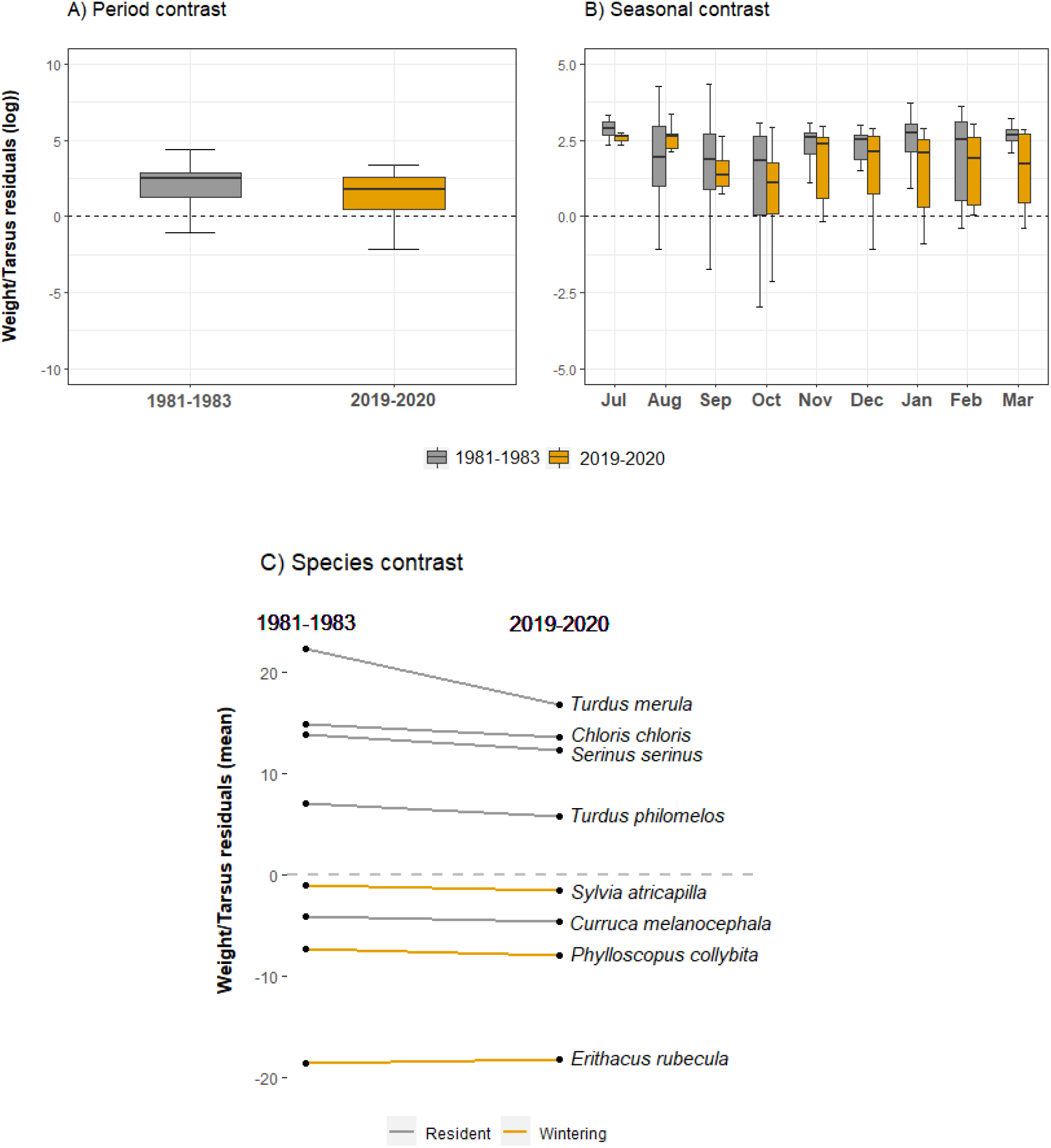
Species body condition change between study periods (A), study months (B) and at species’ level (C). Black (A and B) and grey (C) dotted lines separates positive (“good” body condition) and negative residuals (“poor” body condition). A and B body condition values are expressed in logarithmic scale.

## Discussion

In this study, we explored bird community changes over a 40 year-time span between annual cycles of 1981-1983 and 2019-2020, aiming to test whether avian communities showed significant changes over time in diversity, phenology, morphometry, and body condition. Our results indicate that the avian community is effectively altered in all these aspects, with lasting consequences for ecosystem services and biodiversity conservation.

### Community composition

We have observed a remarkable transformation in Hato Raton’s bird community composition and beta-diversity, with more than half of the main species replaced from 1981-1983. We found out that migrant species are less common in number and abundance, recording a 66 % decrease in wintering species abundance, though we did not detect species turnover in this group, and barely recording transient species in 2019-2020. Together with these losses of migrant species, we also recorded fewer frugivorous and legitimate seed-disperser bird species; resident species that are non-frugivorous and non-disperser were more abundant and rich in number, almost doubling gains in the 2019-2020 records. As migrant birds are likely to find adverse conditions at several points of their migratory routes (Wilcove & Terborgh, 1984), it is difficult to discern the most significant causes of their population decline. Population shifts of European migrant species have been found to be firmly related to climatic conditions on breeding and wintering grounds (such as our study area) and provenance areas, but are also likely to be strongly related to land-use changes (Bellard et al., 2014; Howard et al., 2020). For instance, a warmer climate in northern areas could provide enough resources and a more suitable environment for breeding or even wintering, reducing the need for long-distance flights to find southernmost breeding areas. As species may have started breeding at increasingly higher latitudes, the bird arrival contingent could be reduced in our study area. Moreover, as species differ in their resource tracking capability in shifting scenarios, the more dispersive taxa could be the ones capable of returning to traditional breeding areas (Socolar et al., 2016). Our results are consistent with these interpretations, yet broader-scale data from the Mediterranean Basin would be necessary to test the consistency of these trends in different areas.

Vegetation change in our study area could be another possible explanation for our findings. We detected a considerable increase of forest cover, taller vegetation physiognomies, and a reduction of the cover of fleshy-fruited species, transforming our study area from a shrub-dominated area in the 1980s to an area with larger abundance of pines with poorer understory at present. Transformation of Hato Ratón’s vegetation could directly affect migratory-species habitat selection, as they are commonly found to prefer wide open spaces in edge-type areas in which shrublands with fleshy-fruited species dominate (Zamora et al. 2010). Therefore, spatial distribution of vegetation, changes in plant community structure, and fleshy-fruited plant abundance could explain our diminished detection of migrant birds due to habitat selection, despite breeding ground site fidelity.

Also, we cannot discount the profound transformation that Doñana’s Natural Area has experienced over the last decades. From being a natural space of remarkable plant and animal biodiversity, it has been seriously degraded by agricultural human action. One of the most attractive areas for birds in Doñana, the marshlands, have experienced a 82 % reduction from the beginning of the 20th century to the 2000s decade, mainly as a consequence of human-induced changes, such as desiccation for agricultural land and modification of stream channels (Haberl et al., 2009), both exacerbated by changes in the climatic conditions towards higher temperatures and less predictable precipitation. A final explanation for the observed decrease of migrant species could be the simultaneous enhancement of the resident species local population. As a consequence of area transformation, available resources may not be sufficient to support an avian community as large as in the 1980s. Resident species could prove to be superior competitors, choosing the best feeding areas before migrant species arrive. For instance, de la Hera et al., (2018) found that *Erithacus rubecula*’s resident populations in S Spain tend to occupy woodlands, unlike their migrant conspecifics, which occupy shrublands. This is likely to play a role in the development of local adaptations for the different habitats in the breeding grounds. Similarly, resident individuals of *Sylvia atricapilla* in S Spain are larger in body size and tend to be more abundant in forests (Pérez-Tris & Tellería, 2002), suggesting an advantage over their migrant conspecifics. Overall, vegetation shifts towards greater arboreal cover appear to have favoured a reduced representation of migrants vs. the resident assemblage.

The reduction of legitimate seed-disperser species associated with migrant bird declines could have a major impact on ecosystem function and services (Inger et al., 2014). As interactions are often lost well before species completely disappear (Valiente-Banuet et al., 2015), a reduction of seed dispersers may immediately impact ecosystem functioning, with a potential reduction of plant seed dispersal success and, therefore, direct consequences for the recruitment of fleshy-fruited species and plant community composition. Due to the progressive disappearance of these crucial species, plants whose reproduction relies on them may become less abundant in favour of other plants with abiotic seed dispersal, resulting in a change in plant composition. We have possibly detected this effect in the huge expansion of *Pinus pinea* conifer in our study plots, although this hypothesis warrants further investigation. If this trend is maintained in the coming years, our study area would become increasingly homogeneous, leading to a complete reorganization of the avian community and the fleshy-fruited plants that depend upon it, ultimately leading to an undesirable ecological collapse.

### Bird timing

At a seasonal scale, we found consistent shifts among resident and wintering-species’ seasonal abundance distributions, identifying 13 trends for advancement shifts and a single species, out of a total 15, showing a delay when comparing 1981-1983 vs. in 2019-2020. The proportion of advancement trends versus delays is practically the same within migratory (resident and wintering), trophic (frugivore and non-frugivore) and functional (seed disperser and non-disperser) categories, revealing a consistent and significant shift towards earlier abundance peaks among species.

Due to the lack of continuity between our sampled time points, our results cannot be explained in terms of long-term trends along a temporal series on an annual basis. However, these changes in seasonal abundances are highly relevant and in correspondence with reported temporal advances of migrant species arrival to their wintering grounds as a response to warmer temperatures (Butler, 2003; Haest et al., 2020; Koleček et al., 2020). Migratory behaviour has been widely attributed as a resource tracking mechanism of birds with enough phenotypic plasticity to adapt their timing to shifting food-resource peaks (Haest et al., 2020; van Schaik et al., 1993): whilst plant-feeding species seem to be able to match earlier plant production, birds with an insectivore diet seem to be less headed towards phase advancements (Butler, 2003). This could be the explanation for the lack of seasonal change in abundance found for *P. collybita*. Nevertheless, a complex set of factors could ultimately drive bird phenological advancement: climatic conditions in departure areas could delay departure or adverse weather conditions during migratory flights could affect arrival dates, forcing bird species to increase the number of stopovers or time spent in them during their journey (Gordo, 2007; Moller et al., 2008).

Regarding peak advancement of resident species, this appears to be more related to earlier fruit availability. Advanced plant fructification due to increased temperatures could have induced a continuous advancement of life cycles of resident species over decades. An earlier reproduction and breeding time seem to be necessary to best match juvenile fattening with food availability, which is likely to support the shifts in abundance peaks that we detected. However, species capability to locally adapt to a changing environment is limited. If the current biodiversity change drivers continue, either plant or bird species could reach a limit for their response potential. This means that birds’ food resources may become limited in crucial moments of their cycles. If their fattening is compromised, their abundances may decrease rapidly, which would certainly affect fleshy fruited plants’ dispersal success, limiting seed dispersal in many plant species. In this scenario, a plant community change will likely occur, as potentially already witnessed in our study plots.

### Morphology

We detected long-term changes in the four morphometric parameters. We found significant elongations in tail lengths for *S. atricapilla* and *E. rubecula* (wintering species), in wing length for *S. atricapilla*, and in tarsus length for *C. chloris* (resident), all of them representing ~1.5 % net increase. Many studies have reported morphological changes across several migrant avian taxa in the last decades. Body transformations have two general trends; 1) a decrease in size correlated with temperature rising, which seems to be consistent with the well-known Bergmann’s rule (Andrew et al., 2017; Weeks et al., 2019); and 2), some body components (particularly, wing length) seem elongated in warmer climatic conditions, in line with Allen’s rule (Lank et al., 2017; Nowakowski, 2000; Weeks et al., 2019). However, in other cases, shorter wings have been reported (Van Buskirk et al., 2010). Although these changes are often correlated with increasing temperatures, they have also been hypothesized to cause a change in birds’ shape that may create a more efficient flying body or increased maneuverability (Swaddle & Lockwood, 1998). Our reported ~1.5 % increase over 40 years is in accordance with other researchers’ findings (Gardner et al., 2014; Goodman et al., 2012) revealing that biologically-relevant phenotypic changes may occur very rapidly. Our results partially support an increased wing length in migrant *S. atricapilla*, and, interestingly, we also found an increase in the tail length of two migrant species (*S. atricapilla* and *E. rubecula*) that could also be related with flight efficiency. On the contrary, the tarsus elongation that we detected in resident *C. chloris* is more likely an adaptation to ground-foraging functions, as suggested in other research (Miles & Ricklefs, 1984).

Changes in body size can also be explained by the population composition of measured species, which are able to maintain a settled population in the wintering grounds besides the yearly arrival of the migratory one. Settled species have sometimes been found to have larger body sizes than non-settled, usually also related to a more forestal-like environment (e.g., Pérez-Tris & Tellería, 2002). A reduced arrival of migrant individuals could change the proportion of settled (“resident”) versus migrant conspecifics, pushing higher the mean length values that we recorded. However, this hypothesis is poorly supported by our own data, as we did not record individuals of *S. atricapilla* and *E. rubecula* out of the wintering season in the study area.

### Fat accumulation & body condition index

We identified significant seasonal variation in bird fat accumulation along months and periods. Birds with low fat accumulation are the most common in each included season (approx. 75 % of recorded individuals). On the contrary, medium and high body fat accumulation individuals were less common in the three 2019-2020s migratory seasons than in the 1980s, with their maximum detection delayed from warm months (July-September) to autumn months (October-November). This poorer fat accumulation is in accordance with the trends in body condition index, resulting in a subtle but quantifiable worsening at decadal, seasonal, and specific levels. The detection of a poorer body condition in the months when migrant species should have already acquired their energetic needs to migrate is concerning. Feeding on insects is key to gain a significant body mass in birds. When insect abundance (and therefore, protein resources) in spring or autumn is limited, some migrant species tend to compensate their nutritional intake by switching to a more frugivorous diet (Aamidor et al., 2011; Carnicer et al., 2009). Fruit availability is usually the main determinant of fruit choice in bird species, but it also depends on fruit pulp’s nutrient availability and quantity (Blendinger et al., 2015; Jordano, 1988). However, if fruit availability is drastically reduced, functions like migratory activity, breeding or molt (Bairlein & Gwinner, 1994) may be severely compromised, affecting migratory and reproductive success.

The fact that we have detected an inferior general body condition implies that resident bird species, which are the most abundant in our case, have also experienced this physical deterioration. This is especially apparent in the case of resident *T. merula*, whose decreased body condition in 2019-2020 supports the idea of fruit availability as the main cause of our results. Even though we are confident that fruit production in 1981-1983 was exceptionally high, especially in the 1982-1983 season, the loss of fleshy-fruited species cover in our area (from 72 to 51%) is probably the main cause of this trend. We must consider that, in such a seasonal habitat as our study area, the temporal mismatch between bird species and fruit supply may become exacerbated if global change drivers continue its expansion.

## Conclusive remarks

Bird species have experienced major changes in the last decades that affect almost every aspect of their life cycles, including population dynamics, migratory and feeding behaviour, and morphological traits. Although birds are showing plastic responses to environmental changes, this is unlikely to be sufficient, and the decline signs are increasingly evident. As bird species are globally widespread and largely diverse, identifying the ultimate causes of their changes is an arduous task. However, to focus efforts on ecologically relevant species, as we have done in this manuscript, seems a more attainable objective. It is a priority to diagnose the threats these species are currently facing, as it will help prevent future irreversible effects over Earth’s ecosystems.

## Acknowledgements

This TFM has benefited from the TEMPNET project, funded by a Marie-Sklodowska Curie Fellowship (798269 - TEMPNET - H2020-MSCA-IF-2017) by the European Commission to IM, and in part with funding from project CGL 2017-82847 from Spanish Ministry of Science and Innovation (PJ). Data from the 80s come from the PhD Thesis work of Pedro Jordano. Carlos Gutiérrez-Expósito (CGE) and Julio Rabadán González (JRG) were in charge of bird monitoring and ringing in 2019-2020. JA Sarrión, J. Rengel and a dozens of volunteers also helped in bird trapping. Amy Dennett and Rafe Cotton assisted with field work. The help from staff and guard parks of the Doñana National Park and the ICTS-RBD providing access authorizations and logistic support is highly appreciated. Ana Benítez López (ABL) has played a pivotal role in this TFM preparation. James Martin helped with the language edit of the text.

## Author contribution

IM, ABL and PJ conceived the original idea. IM, PJ, JRG and CGE collected data. MCC collated historical data from the 80s, performed data analyses and led manuscript writing. All authors critically commented on previous drafts and edited the final version of the manuscript.

