## Supplementary Material for "Assessing short- and long-term variations in diversity, timing, and body condition of migratory frugivorous birds"

### Tables:

**Table S1:** Vegetation cover comparison between year groups. “Per transect” contains transects mean and confidence interval (“IC”) or standard error (“SE”). “Other spp.” are collapsed categories of the less abundant plant species. “Fleshy fruit” is a collapsed category of fleshy-fruited species. “% of covered area” is the percentage of area covered by any type of vegetation, i.e. total area minus percentage of bare ground. “Presence” column indicates the percentage of transects in which a species was found (mean in “Other spp.”).

| Year group | Species | Fruit type | Total % | Per transect | | Presence |
|---|---|---|---|---|---|---|
| | | | | Mean | IC | |
| 1981 - 1983 | *Pistacia lentiscus* | Fleshy fruit | 33.40 | 35.00 | 11.30 | 100.00 |
| | *Smilax aspera* | Fleshy fruit | 12.40 | 13.40 | 12.60 | 86.30 |
| | *Ulex parviflorus* | Other | 8.60 | 6.60 | 9.70 | 40.00 |
| | *Olea europaea* | Fleshy fruit | 7.80 | 8.00 | 8.50 | 73.30 |
| | *Halimium halimifolium* | Other | 4.90 | 3.10 | 6.80 | 26.70 |
| | *Phillyrea angustifolia* | Fleshy fruit | 4.80 | 4.70 | 4.60 | 86.70 |
| | *Cistus salvifolius* | Other | 4.30 | 3.40 | 5.60 | 60.00 |
| | *Rhamnus lycioides* | Fleshy fruit | 3.90 | 4.50 | 4.70 | 80.00 |
| | *Ruscus aculeatus* | Fleshy fruit | 2.40 | 2.40 | 3.70 | 60.00 |
| | *Rubus ulmifolius* | Fleshy fruit | 1.90 | 2.10 | 4.50 | 26.70 |
| | **Other 12 spp.** | Fleshy fruit/Other | 7.52 | 0.79 | 0.17 | 25.00 |
| | **Fleshy fruit (16 spp.)** | | **72.11** | **4.80** | **4.07** | **47.89** |
| | **Bare ground** | | **7.90** | **5.30** | **7.80** | - |
| | **% of covered area** | | **92.10** | - | - | - |

| Year group | Species | Fruit type | Total % | Per transect | | Presence |
|---|---|---|---|---|---|---|
| | | | | Mean | SE | |
| 2019 - 2020 | *Pistacia lentiscus* | Fleshy fruit | 26.65 | 27.40 | 4.56 | 93.33 |
| | *Pinus pinea* | Other | 21.38 | 34.54 | 5.49 | 53.33 |
| | *Cistus salvifolius* | Other | 12.45 | 17.59 | 4.20 | 86.67 |
| | *Ulex parviflorus* | Other | 8.37 | 11.52 | 2.67 | 80.00 |
| | *Phillyrea angustifolia* | Fleshy fruit | 8.04 | 10.35 | 1.76 | 73.33 |
| | *Olea europaea* | Fleshy fruit | 6.87 | 10.64 | 2.57 | 66.67 |
| | *Lavandula stoechas* | Other | 3.57 | 10.76 | 3.02 | 40.00 |
| | *Rubia peregrina* | Fleshy fruit | 2.63 | 4.40 | 1.96 | 40.00 |
| | *Rhamnus lycioides* | Fleshy fruit | 2.57 | 4.93 | 1.50 | 46.67 |
| | *Halimium calycinum* | Other | 1.58 | 6.39 | 2.16 | 33.33 |
| | **Other 15 spp.** | Fleshy fruit/Other | 5.90 | 2.64 | 1.39 | 17.33 |
| | **Fleshy fruit (15 spp.)** | | **50.50** | **3.37** | **1.41** | **33.33** |
| | **Bare ground** | | **27.49** | **31.72** | **5.42** | **86.67** |
| | **% of covered area** | | **72.51** | - | - | - |

**Table S2:** Mean, confidence interval ("Int.") and maximum and minimum registered values of monthly mean temperatures ("Mean T"), monthly minimum temperatures ("Min T") and accumulated rainfall. Part A of the table shows decadal values (in red for study periods) and part B shows specific study years. We obtained monthly total precipitation and minimum and maximum temperature data for the study site from 1981 to 2020 from the CSIC's weather station placed in Doñana Biological Reserve, less than 20 km away from our study site.

| | Years | Mean T (ºC) | | | | Min T (ºC) | | | | Rainfall (mm) | | | |
|---|---|---|---|---|---|---|---|---|---|---|---|---|---|
| | | Mean | Int. | Max. | Min. | Mean | Int. | Max. | Min. | Mean | Int. | Max. | Min. |
| A | 1980-1990* | 16.72 | ± 0.65 | 18.68 | 15.38 | 4.7 | ± 0.98 | 17 | -6 | 547.24 | ± 151.97 | 391.5 | 0 |
| A | 1990-2000 | 16.98 | ± 0.33 | 17.83 | 16.18 | 5.4 | ± 0.67 | 14 | -4.5 | 541.48 | ± 160.71 | 425.9 | 0 |
| A | 2000-2010 | 17.24 | ± 0.28 | 18.24 | 16.58 | 5.98 | ± 0.63 | 15 | -5 | 574.02 | ± 112.97 | 261 | 0 |
| A | 2010-2020* | 17.58 | ± 0.29 | 18.36 | 16.95 | 5.41 | ± 0.49 | 15 | -6 | 496.57 | ± 76.31 | 222.81 | 0 |
| B | 1981-1983 | 16.1 | ± 1.03 | 23.9 | 8.05 | 3.81 | ± 1.04 | 11 | -6 | 342.4 | ± 131.62 | 155.5 | 0 |
| B | 2019-2020 | 17.18 | ± 1.28 | 25.37 | 9.98 | 5.38 | ± 0.73 | 15 | -2 | 432.25 | ± 9.7 | 209.8 | 0 |

**Table S3:** complete list of identified species during censuses and netting sampling, including English and Spanish common names, migration type and subtype, and also trophic and functional categories (113 spp.) From 1 to 6, the ones selected as “representative species”. From 1 to 27, the 20 most abundant ones of both study periods.

| Nº | Scientific name | Common name | Spanish common name | Migration type | Subtype | Trophic | Functional |
|---|---|---|---|---|---|---|---|
| 1 | ***Chloris chloris*** | European greenfinch | Verderón europeo | Resident | European | Granivore-Frugivore | SP |
| 2 | ***Curruca melanocephala*** | Sardinian warbler | Curruca cabecinegra | Resident | | Frugivore-Insectivore | SD |
| 3 | ***Erithacus rubecula*** | European robin | Petirrojo | Wintering | European | Frugivore-Insectivore | SD |
| 4 | ***Phylloscopus collybita*** | Common chiffchaff | Mosquitero común | Wintering | European | Insectivore | NF |
| 5 | ***Sylvia atricapilla*** | Eurasian blackcap | Curruca capirotada | Wintering | European | Frugivore-Insectivore | SD |
| 6 | ***Turdus merula*** | Common blackbird | Mirlo común | Resident | | Frugivore-Insectivore | SD |
| **End of representative species' list** | | | | | | | |
| 7 | ***Aegithalos caudatus*** | Long-tailed tit | Mito | Resident | | Insectivore-Frugivore | PC |
| 8 | ***Carduelis carduelis*** | European goldfinch | Jilguero europeo | Resident | European | Granivore | SP |
| 9 | ***Coloeus monedula*** | Western jackdaw | Grajilla occidental | Resident | | Omnivore | SD |
| 10 | ***Columba palumbus*** | Common wood pigeon | Paloma torcaz | Resident | | Herbivore | SD |
| 11 | ***Curruca undata*** | Dartford warbler | Curruca rabilarga | Resident | | Insectivore-Frugivore | SD |
| 12 | ***Cyanopica cooki*** | Iberian magpie | Rabilargo | Resident | | Omnivore | SD |
| 13 | ***Ficedula hypoleuca*** | European pied flycatcher | Papamoscas cerrojillo | Transient | Trans Saharian | Insectivore-Frugivore | SD |
| 14 | ***Fringilla coelebs*** | Common chaffinch | Pinzón vulgar | Resident | European | Insectivore-Granivore | PC/SP/SD |
| 15 | ***Galerida cristata*** | Crested lark | Cogujada común | Resident | European | Insectivore-Granivore | NF |
| 16 | ***Lanius meridionalis*** | Iberian grey shrike | Alcaudón real | Resident | | Carnivore-Frugivore | SD |
| 17 | ***Lanius senator*** | Woodchat shrike | Alcaudón común | Summering | Trans Saharian | Carnivore-Frugivore | SD |
| 18 | ***Linaria cannabina*** | Common linnet | Pardillo común | Resident | European | Insectivore-Granivore | NF |
| 19 | ***Miliaria calandra*** | Corn bunting | Escribano triguero | Resident | | Granivore | NF |
| 20 | ***Parus major*** | Great tit | Carbonero común | Resident | | Insectivore-Frugivore | PC/SP/SD |
| **Nº** | **Scientific name** | **Common name** | **Spanish common name** | **Migration type** | **Subtype** | **Trophic** | **Functional** |
| 22 | ***Saxicola rubicola*** | European stonechat | Tarabilla europea | Resident | | Insectivore-Frugivore | SD |
| 23 | ***Serinus serinus*** | European serin | Verdecillo | Resident | European | Granivore | PC/SP |
| 24 | ***Sturnus unicolor*** | Spotless starling | Estornino negro | Resident | | Insectivore-Frugivore | SD |
| 25 | ***Sylvia borin*** | Garden warbler | Curruca mosquitera | Transient | Trans Saharian | Frugivore-Insectivore | SD |
| 26 | ***Troglodytes troglodytes*** | Eurasian wren | Chochín común | Resident | | Insectivore | NF |

| | | | | | | | |
|---|---|---|---|---|---|---|---|
| 27 | ***Turdus philomelos*** | Song thrush | Zorzal común | Wintering | European | Frugivore-Insectivore | SD |
| | | | **End of 20 most abundant species' list of each study period** | | | | |
| 28 | ***Accipiter nisus*** | Eurasian sparrowhawk | Gavilán común | Resident | European | Carnivore | NF |
| 29 | ***Acrocephalus scirpaceus*** | Eurasian reed warbler | Carricero común | Summering | Trans Saharian | Insectivore | NF |
| 30 | ***Alauda arvensis*** | Eurasian skylark | Alondra común | Wintering | Resident | Granivore-Insectivore | NF |
| 31 | ***Alcedo atthis*** | Common kingfisher | Martín pescador | Resident | | Carnivore | NF |
| 32 | ***Alectoris rufa*** | Red-legged partridge | Perdiz roja | Resident | | Herbivore | NF |
| 33 | ***Anas platyrhynchos*** | Mallard | Ánade azulón | Resident | | Herbivore | NF |
| 34 | ***Anser anser*** | Greylag goose | Ánsar común | Wintering | European | Herbivore | NF |
| 35 | ***Anthus pratensis*** | Meadow pipit | Bisbita pratense | Wintering | European | Insectivore | NF |
| 36 | ***Apus apus*** | Common swift | Vencejo común | Summering | Trans Saharian | Insectivore | NF |
| 37 | ***Apus pallidus*** | Pallid swift | Vencejo pálido | Summering | Trans Saharian | Insectivore | NF |
| 38 | ***Ardea cinerea*** | Grey heron | Garza real | Resident | | Carnivore | NF |
| 39 | ***Asio otus*** | Long-eared owl | Búho chico | Resident | | Carnivore | NF |
| 40 | ***Athene noctua*** | Little owl | Mochuelo europeo | Resident | | Carnivore | NF |
| 41 | ***Bubo bubo*** | Eurasian eagle-owl | Búho real | Resident | | Carnivore | NF |
| 42 | ***Burhinus oedicnemus*** | Eurasian stone-curlew | Alcaraván común | Resident | European | Omnivore | NF |
| 43 | ***Buteo buteo*** | Common buzzard | Busardo ratonero | Resident | | Carnivore | NF |
| 44 | ***Caprimulgus europaeus*** | European nightjar | Chotacabras europeo | Transient | Trans Saharian | Insectivore | NF |
| 45 | ***Caprimulgus ruficollis*** | Red-necked nightjar | Chotacabras cuellirrojo | Summering | Trans Saharian | Insectivore | NF |
| 46 | ***Cecropis daurica*** | Red-rumped swallow | Golondrina dáurica | Summering | Trans Saharian | Insectivore | NF |
| 47 | ***Certhia brachydactyla*** | Short-toed treecreeper | Agateador común | Resident | | Insectivore | NF |
| **Nº** | **Scientific name** | **Common name** | **Spanish common name** | **Migration type** | **Subtype** | **Trophic** | **Functional** |
| 49 | ***Ciconia ciconia*** | White stork | Cigüeña blanca | Resident | Trans Saharian | Omnivore | NF |
| 50 | ***Ciconia nigra*** | Black stork | Cigüeña negra | Wintering | Trans Saharian | Omnivore | NF |
| 51 | ***Circaetus gallicus*** | Short-toed snake eagle | Águila culebrera | Summering | Trans Saharian | Carnivore | NF |
| 52 | ***Circus aeruginosus*** | Western marsh harrier | Aguilucho lagunero | Resident | Resident | Carnivore | NF |
| 53 | ***Circus cyaneus*** | Hen harrier | Aguilucho pálido | Resident | | Carnivore | NF |

| | | | | | | | |
|---|---|---|---|---|---|---|---|
| 54 | ***Cisticola juncidis*** | Zitting cisticola | Buitrón | Resident | | Insectivore | NF |
| 55 | ***Coccothraustes coccothraustes*** | Hawfinch | Picogordo común | Resident | European | Granivore-Frugivore | SP |
| 56 | ***Columba oenas*** | Stock dove | Paloma zurita | Wintering | Resident | Herbivore | NF |
| 57 | ***Corvus corax*** | Common raven | Cuervo grande | Resident | | Omnivore | SD |
| 58 | ***Corvus corone*** | Carrion crow | Corneja común | Resident | | Omnivore | SD |
| 59 | ***Curruca cantillans*** | Western subalpine warbler | Curruca carrasqueña | Transient | Trans Saharian | Frugivore-Insectivore | SD |
| 60 | ***Curruca communis*** | Common whitethroat | Curruca zarcera | Transient | Trans Saharian | Frugivore-Insectivore | SD |
| 61 | ***Curruca conspicillata*** | Spectacled warbler | Curruca tomillera | Summering | Resident | Insectivore-Frugivore | SD |
| 62 | ***Curruca hortensis*** | Western Orphean warbler | Curruca mirlona | Transient | Trans Saharian | Frugivore-Insectivore | SD |
| 63 | ***Cyanistes caeruleus*** | Eurasian blue tit | Herrerillo común | Resident | | Insectivore-Frugivore | PC/SP/SD |
| 64 | ***Delichon urbicum*** | Common house martin | Avión común | Summering | Trans Saharian | Insectivore | NF |
| 65 | ***Dendrocopos major*** | Great spotted woodpecker | Pico picapinos | Resident | | Omnivore | PC/SP/SD |
| 66 | ***Elanus caeruleus*** | Black-winged kite | Elanio común | Resident | | Carnivore | NF |
| 67 | ***Estrilda astrild*** | Common waxbill | Estrilda común | Resident | | Granivore | SP |
| 68 | ***Falco tinnunculus*** | Common kestrel | Cernícalo vulgar | Resident | European / Trans Saharian | Carnivore | NF |
| 69 | ***Galerida theklae*** | Thekla's lark | Cogujada montesina | Resident | | Insectivore-Granivore | NF |
| 70 | ***Gallinula chloropus*** | Common moorhen | Gallineta común | Resident | | Omnivore | NF |
| 71 | ***Grus grus*** | Common crane | Grulla común | Wintering | European | Herbivore-Insectivore | NF |
| 72 | ***Gyps fulvus*** | Griffon vulture | Buitre leonado | Resident | | Necrophagous | NF |
| 73 | ***Hieraaetus pennatus*** | Booted eagle | Águila calzada | Summering | Trans Saharian | Carnivore | NF |
| 74 | ***Hippolais polyglotta*** | Melodious warbler | Zarcero políglota | Summering | Trans Saharian | Insectivore | NF |
| **Nº** | **Scientific name** | **Common name** | **Spanish common name** | **Migration type** | **Subtype** | **Trophic** | **Functional** |
| 76 | ***Iduna pallida*** | Eastern olivaceous warbler | Zarcero pálido | | | | |
| 77 | ***Jynx torquilla*** | Eurasian wryneck | Torcecuellos euroasiático | Summering | Resident / Trans Saharian | Insectivore | NF |
| 78 | ***Larus fuscus*** | Lesser black-backed gull | Gaviota sombría | Wintering | European | Omnivore | NF |
| 79 | ***Lophophanes cristatus*** | European crested tit | Herrerillo capuchino | Resident | | Insectivore-Frugivore | PC |
| 80 | ***Lullula arborea*** | Woodlark | Alondra totovía | Resident | | Insectivore-Granivore | NF |

| | | | | | | | |
|---|---|---|---|---|---|---|---|
| 81 | ***Luscinia megarhynchos*** | Common nightingale | Ruiseñor común | Summering | Trans Saharian | Insectivore-Frugivore | SD |
| 82 | ***Merops apiaster*** | European bee-eater | Abejaruco europeo | Summering | Trans Saharian | Insectivore | NF |
| 83 | ***Milvus migrans*** | Black kite | Milano negro | Summering | Trans Saharian | Carnivore | NF |
| 84 | ***Milvus milvus*** | Red kite | Milano real | Resident | European | Carnivore | NF |
| 85 | ***Motacilla alba*** | White wagtail | Lavandera blanca | Wintering | Resident | Insectivore | NF |
| 86 | ***Motacilla cinerea*** | Grey wagtail | Lavandera cascadeña | Resident | European | Insectivore | NF |
| 87 | ***Motacilla flava*** | Western yellow wagtail | Lavandera boyera | Summering | European | Insectivore | NF |
| 88 | ***Muscicapa striata*** | Spotted flycatcher | Papamoscas gris | Summering | Trans Saharian | Insectivore-Frugivore | SD |
| 89 | ***Oriolus oriolus*** | Eurasian golden oriole | Oropéndola europea | Summering | Trans Saharian | Insectivore-Frugivore | SD |
| 90 | ***Passer domesticus*** | House sparrow | Gorrión común | Resident | | Granivore-Insectivore | PC/SP |
| 91 | ***Passer hispaniolensis*** | Spanish sparrow | Gorrión moruno | Resident | | Granivore-Insectivore | PC/SP |
| 92 | ***Phalacrocorax carbo*** | Great cormorant | Cormorán grande | Wintering | Resident | Carnivore | NF |
| 93 | ***Phoenicurus ochruros*** | Black redstart | Colirrojo tizón | Wintering | European | Insectivore-Frugivore | SD |
| 94 | ***Phoenicurus phoenicurus*** | Common redstart | Colirrojo real | Transient | Trans Saharian | Insectivore-Frugivore | SD |
| 95 | ***Phylloscopus bonelli*** | Western Bonelli's warbler | Mosquitero papialbo | Transient | Trans Saharian | Insectivore | NF |
| 96 | ***Phylloscopus ibericus*** | Iberian chiffchaff | Mosquitero ibérico | Summering | Trans Saharian | Insectivore | NF |
| 97 | ***Phylloscopus trochilus*** | Willow warbler | Mosquitero musical | Transient | Trans Saharian | Insectivore | NF |
| 98 | ***Picus sharpei*** | Iberian green woodpecker | Pito ibérico | Resident | | Insectivore | NF |
| 99 | ***Platalea leucorodia*** | Eurasian spoonbill | Espátula común | Resident | | Carnivore | NF |
| 100 | ***Porphyrio porphyrio*** | Western swamphen | Calamón común | Resident | | Herbivore | NF |
| 101 | ***Prunella modularis*** | Dunnock | Acentor común | Wintering | European | Insectivore-Frugivore | SD |
| **Nº** | **Scientific name** | **Common name** | **Spanish common name** | **Migration type** | **Subtype** | **Trophic** | **Functional** |
| 102 | ***Pyrrhula pyrrhula*** | Eurasian bullfinch | Camachuelo común | Wintering | European | Herbivore-Frugivore | PC/SP |
| 103 | ***Regulus ignicapilla*** | Common firecrest | Reyezuelo listado | Resident | European | Insectivore | NF |
| 104 | ***Remiz pendulinus*** | Eurasian penduline tit | Pájaro moscón | Resident | | Insectivore | NF |
| 105 | ***Spinus spinus*** | Eurasian siskin | Jilguero lúgano | Wintering | European | Granivore | NF |
| 106 | ***Streptopelia decaocto*** | Eurasian collared dove | Tórtola turca | Resident | | Herbivore | NF |
| 107 | ***Streptopelia turtur*** | European turtle dove | Tórtola europea | Summering | Trans Saharian | Herbivore | NF |

| | | | | | | | |
|---|---|---|---|---|---|---|---|
| 108 | ***Strix aluco*** | Tawny owl | Cárabo común | Resident | | Carnivore | NF |
| 109 | ***Tachybaptus ruficollis*** | Little grebe | Zampullín común | Resident | | Insectivore-Carnivore | NF |
| 110 | ***Tringa ochropus*** | Green sandpiper | Andarríos grande | Wintering | European | Insectivore | NF |
| 111 | ***Turdus iliacus*** | Redwing | Zorzal alirrojo | Wintering | European | Frugivore-Insectivore | SD |
| 112 | ***Turdus viscivorus*** | Mistle thrush | Zorzal charlo | Resident | European | Frugivore-Insectivore | SD |
| 113 | ***Upupa epops*** | Eurasian hoopoe | Abubilla | Resident | Trans Saharian | Insectivore | NF |

**Figures:**

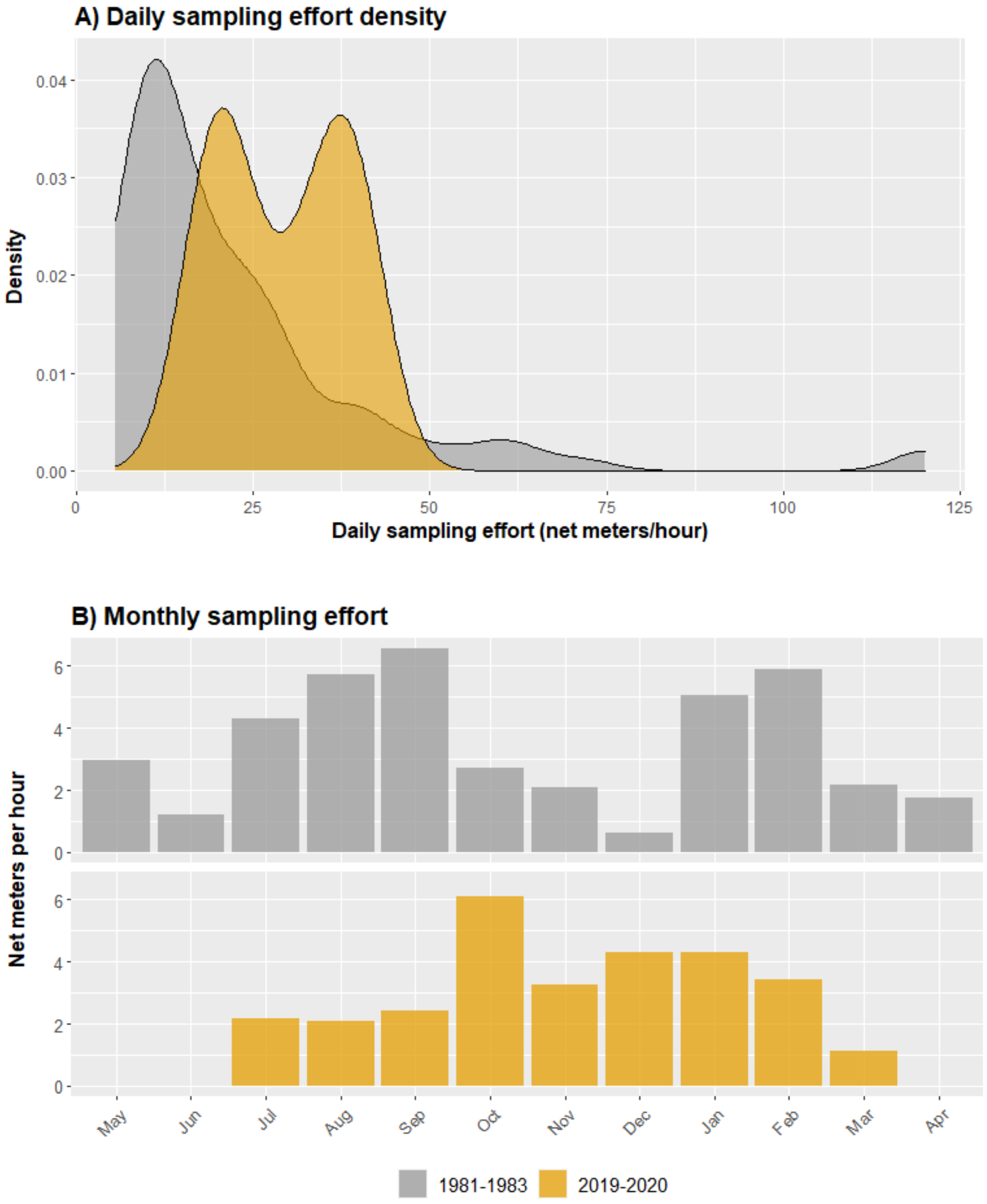


**Figure S1:** Birds' netting daily sampling effort density (A) and monthly sampling effort (B). Although in 2019-2020, the daily number of net meters operating per hour was higher (25-50 m/h) than in 1981-1983 (10-25 m/h), 1981-1983 period accumulates much more net meters/hour per month, due to higher sampling frequency (weekly in 1981-1983, biweekly in 2019-2020).

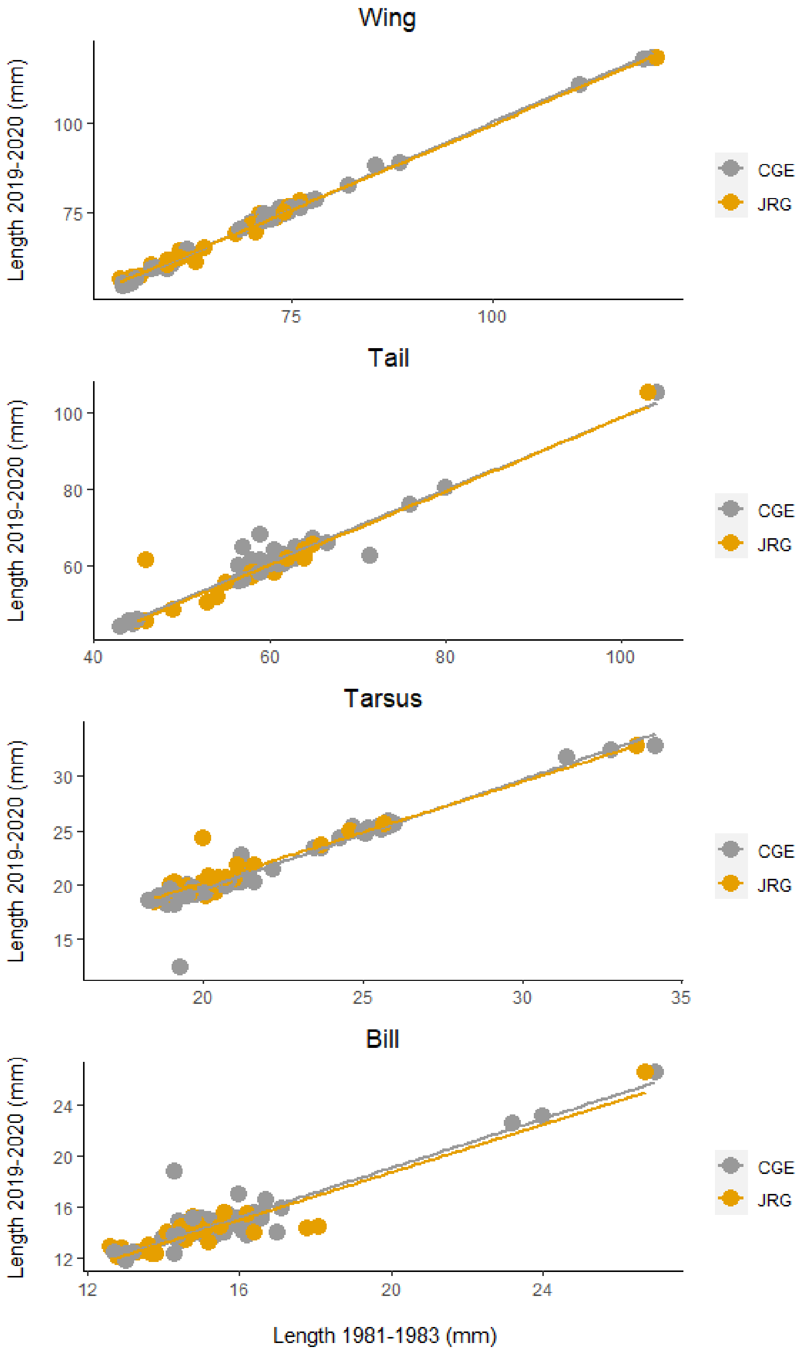


**Figure S2:** Repeatability of sampling measures across researchers. The same individuals were measured in 2019-2020 by Pedro Jordano (the only ringer in the 80s, values belonging to x-axis) and the other two ringers (CGE or JRG, values represented in y-axis). These scatterplots show the correlation among measures, being the correlation coefficient higher than 0.92 in all variables.

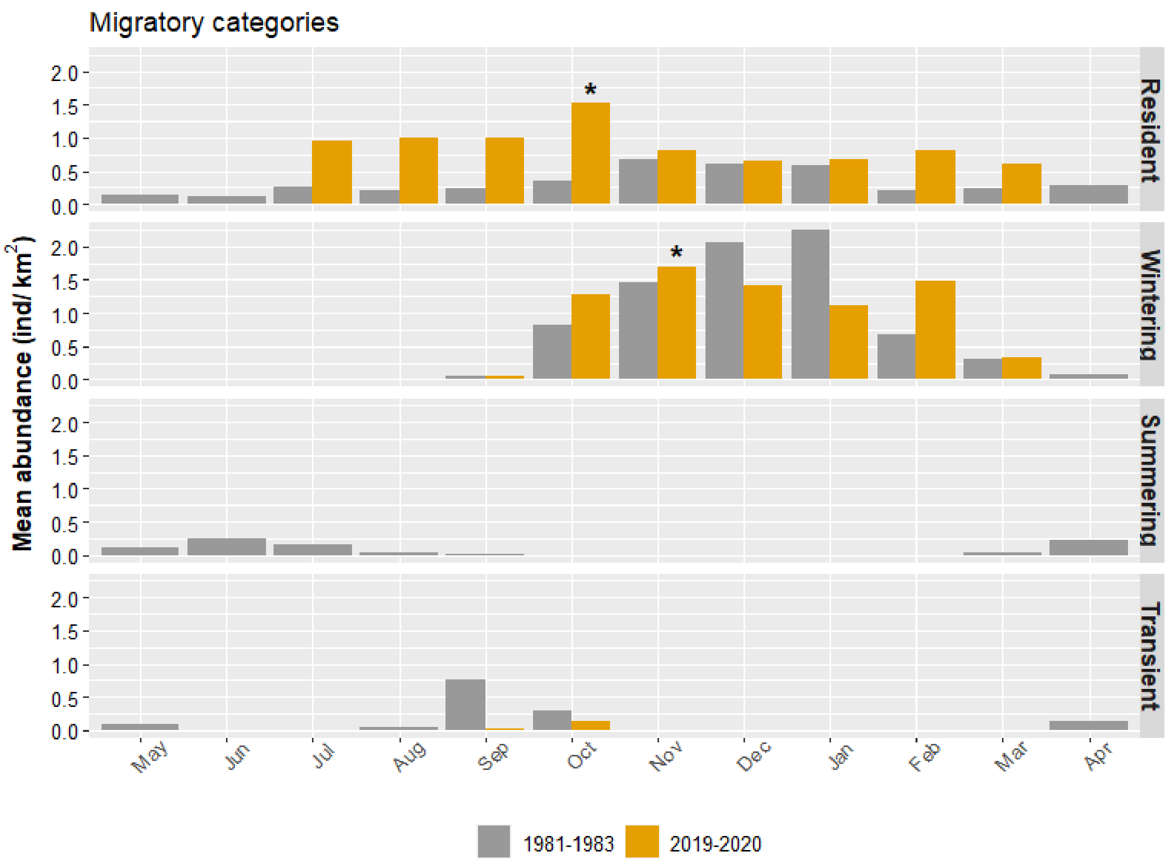


**Figure S3:** Phenological bar plots of monthly mean densities (individuals per km²) in collapsed species' migratory groups: resident, wintering, summering, and transient bird species. In grey colour the 1981-1983 recordings and, in orange, the 2019-2020 ones. Black asterisks mark the main abundance peak in the 2019-2020 period.

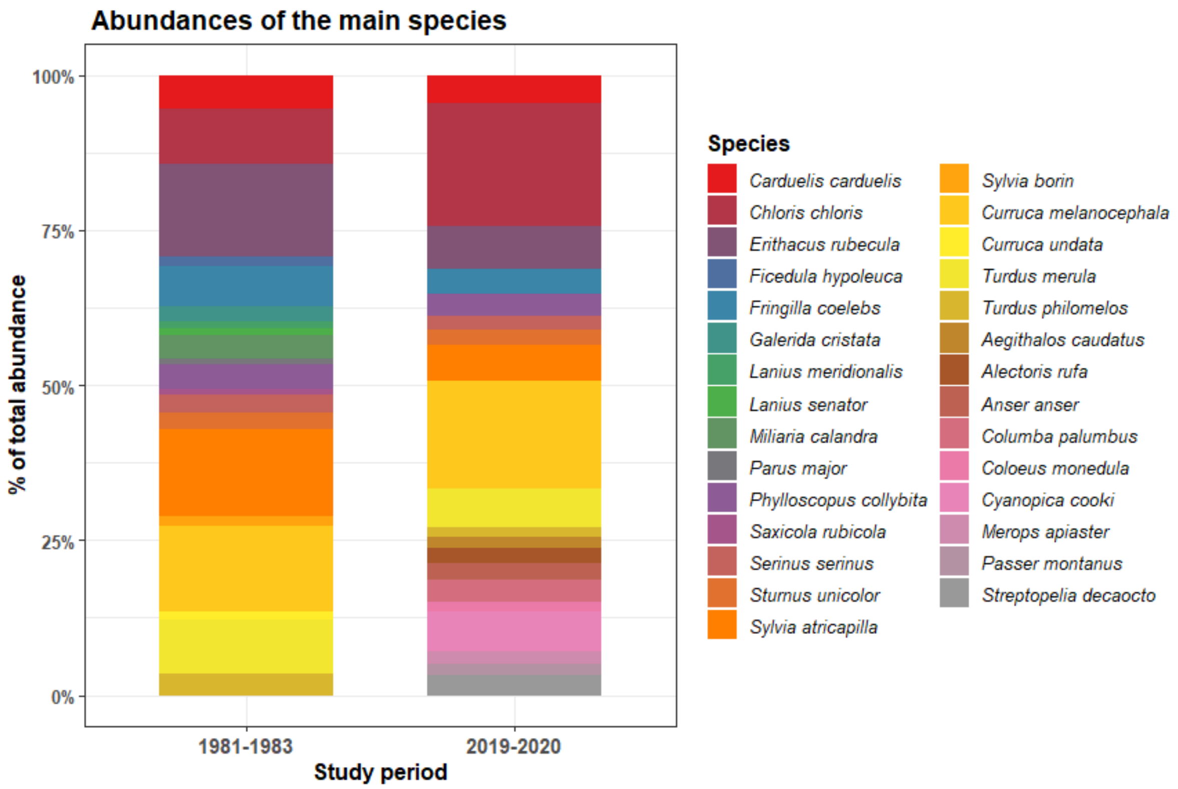


**Figure S4:** Species abundance contrast between time periods. Relation of the 20 most recorded species in each studied period of time. The height of the coloured blocks is proportional to the percentage of species abundance.
